# Effects of polyploidy on response of *Dunaliella salina* to salinity

**DOI:** 10.1101/219840

**Authors:** Fatemeh Soltani Nezhad, Hakimeh Mansouri

**Affiliations:** Department of Biology, Faculty of Science, Shahid Bahonar University of Kerman, Kerman, Iran

**Keywords:** Antioxidant Enzymes, Carotenoid, *Dunaliella salina*, Glycerol, Polyploidy

## Abstract

In this study, effect of different percentages of polyploid cells of *Dunaliella salina* in culture medium, on growth and other biochemical parameters of algae under different salinity levels were investigated. The results indicated that concentration 3M NaCl is the optimal concentration of salt, since in this concentration, the highest enhancement in fresh and dry weight, chlorophyll and carotenoids, soluble sugar, glycerol, protein and starch content was observed in comparison with other concentrations. The amount of these metabolites declined in the concentrations under the optimum salinity. The least and highest amounts of MDA were observed at 1 and 4 M NaCl respectively. Polyploidy in optimum concentration of salt, caused further increment of the above growth parameters. In this relation, in most cases, treatment of 0.1% was more effective. The beneficial effects of polyploidy in non-optimal conditions were also found in some parameters such as biomass, chlorophyll, carotenoids, proteins and starch. The activity of antioxidant enzymes CAT, SOD and POD were maximum in 4 M NaCl. Polyploidy affected activity of these antioxidant enzymes in some concentrations of salt. Overall, our results suggest that the microalgae have significantly different responses to salt stress based on ploidy level of the algae.

**Abbreviations:**

AOS: Active Oxygen Products
CAT: Catalase
GPX: Guaiacol Peroxidase
EDTA: Ethylene Diamine Tetraacetic Acid
MDA: Malondialdehyde
PMSF: Phenyl Methanesulfonyl Fluoride
PVP: Polyvinylpyrrolidone
SOD: Superoxide Dismutase

## 1. Introduction

Salinity stress is considered one of the most significant abiotic stresses, affecting various aspects of metabolism and physiology. In order to understand how algae respond and adapt to salinity changes, the biochemical and physiological responses were thoroughly studied (Kirst, 1990). Anyway, information on the effects of salinity stress on the induction of oxidative stress is low in algae (Jahnke and White, 2003; Liu *et al.,* 2007). Adapting to salinity in various ways, Algae are divided into two groups according to their tolerance as halophytic (salt requiring for optimum growth) and halo tolerant (having response mechanisms allowing them to live in saline environments) (Rao *et al.,* 2007). As a unicellular photoautotroph halo-tolerant green algae, *Dunaliella* lacks a rigid cell wall. Furthermore, it can store many economically important organic compounds such as β-carotene, glycerol and others (Ghetti *et al.,* 1999). Additionally, *Dunaliella* responds to salt stress by high accumulation of glycerol and certain proteins as well as enhanced elimination of Na^+^ ions (Pick, 2002). However, the function and physiological role of glycerol may be different in each *Dunaliella* species. Moreover, it seems to function as an osmotic regulator such that cells without wall should maintain iso-osmotic conditions with their environment (Hadi *et al.,* 2008). Owing to changing external salinity, the basic biochemical response is observed at internal glycerol level, changing the external salinity directly (Ben-Amotz, 1975, Chitlaru and Pick, 1991). Carotenoids (astaxanthin and lutein) act as complementary pigments in the harvesting complex and as protective agents against the active oxygen products (AOS) formed from photo-oxidation. They are also an integral part of the photosynthetic apparatus in the algae. These oxygen radicals can react with macromolecules and lead to cellular damages (Malanga *et al.,* 1997). The algae have developed defensive systems against photo-oxidative damage and can remove these highly reactive oxygen species by antioxidative mechanisms to detoxify. These antioxidant defensive system consist of hydrophobic (carotenoids and α-tocopherol) and hydrophilic antioxidant (ascorbic acid and glutathione) and antioxidant enzymes such as superoxide dismutase (SOD), catalsae (CAT), glutathione reductase (GR) ascorbic peroxidase (APX) and peroxidase (POD) (Rao *et al.,* 1996; Malanga *et al.,* 1997; Rijstenbil, 2002). In previous reports, it was mentioned that *D. salina* elevated the activities of the antioxidant enzymes and accumulated a large amount of carotenoids, α-tocopherol and ascorbic acid when grown in a culture medium containing high salt concentration and/or limiting nitrogen (El-Baz *et al.,* 2002).

Due to the interest in a homogenous genetic background, some studies have conducted in recent decades to compare polyploids and their diploid relatives for tolerance to environmental stressors. Polyploidy is caused by doubling chromosomes of a single species (autopolyploidy) or the hybrids between two species (allopolyploidy). Polyploids clearly show higher resistance to drought, heat, cold, salt, viruses, fungi, and pest stressors compared with their diploid relatives Zhang *et al.,* 2010). Therefore, polyploidy has been considered an effective way to increase the resistance to environmental stresses in plants, playing important roles in agriculture and forestry (Xiong and Zhang, 2006). Polyploidy also varies in physiological functions or gene expressions. Accordingly, these alterations might affect the response to stress and many phenotypes (Wang *et al.,* 2013). Comparative cytochemical studies of untreated and colchicine-treated cells of algae (*Chlamydomonas*) illustrated an enhancement in the content of starch, protein, and lipid materials. The data also suggest that the enlarged walls of treated cells are composed of a complex polysaccharide (Walne, 1966). The aim of this work was to study the effect of polyploidy on many important metabolites and antioxidant enzymatic system of *D. salina* when grown in the medium containing different levels of NaCl.

## 2. MATERIALS AND METHODS

### 2.1. Growth conditions

*D. salina* was isolated from the Salt River of Shahdad (N30°24′16.164′′, E57^°^40′57.828′′) in Kerman, Iran and identified based on physiological and morphological descriptions in the references mentioned by Massyuk, (1973c) as well as Browitzka and Siva, (2007). After transferring the samples into the laboratory, EC (33.5 ds cm^-1^) and pH (7.75) of water were measured. Algae were cultured on the agar plate. After 2 to 3 weeks, each colony was transferred to 20 ml liquid growth medium (Artificial Sea Water, ASW) containing 50 mM NaHCO_3_, 5 mM MgSO_4_, 0.75 mM KNO_3_, 0.2 mM KH_2_PO_4_, 0.2 mM CaCl_2_, 7 μM MnCl_2_, 5 μM EDTA, 2 μM FeCl_3_, 1 μM CuCl_2_, 1μM CoCl_2_, 1 μM (NH_4_) 6 Mo_7_O_24_, and 1 μM ZnCl_2_ with NaCl added to obtain the required salinity medium at 2 M (pH 7.5). The cultures were incubated in a growth chamber under 16/8 h light-dark provided by cool white fluorescent lamps at the intensity of 49 μmol photons m^−^2 s^−^1 at 25 ± 2 °C, and were shaken manually twice a day to ensure a uniform illumination of the cells and stock cultures were sub-cultured at least once per month.

### 2.2. Polyploidy induction by colchicine

Haploid vegetative cells of *D. salina* were exposed to concentrations of 0.1 and 0.5% colchicine (Sigma-Aldrich), made in culture solution for 36 hours. After passing these times, algae cells were washed completely for two or three times with culture solution to free algae from traces of the alkaloid (Sarma, 1957) and then were finally transferred to a sterile fresh medium for 21 days. Afterward, the treated cells (0.5×10^3^ cells per μl) with colchicine were counted by cell counter. The fresh cultures containing 1, 2, 3 and 4 molar NaCl with 3 replications per treatment were provided and inoculated with 10 ml (5×10^6^ cells) stock cultures of every colchicine treatment (0, 0.1 and 0.5 %). The various treated samples were allowed to grow under the same conditions as control samples. At the end of three weeks, all samples were centrifuged and the pellets obtained were then frozen and stored at −70 °C prior to analysis.

### 2.3. Flow cytometry analysis

Ploidy level of the cells of *D. salina* treated with different colchicine concentrations at different time of exposure was determined by BD FACSCalibur cell sorting System system flow cytometry (FCM) (USA) equipped with two lasers. First, the cells were harvested by centrifuging at 3000 rpm for 5 minutes. After discarding the supernatant, the obtained pellet was resuspended in a specific lysis buffer (300μl) of kit related to flow cytometry device with high resolution DNA analysis of nuclei optimal for some plant species and microorganisms. This kit has two steps: disaggregation and propidium iodide staining. Nuclear suspensions were filtered through a 50 μm nylon filter and 5 μl of propidium idoid 0.1 mg mL^-1^ (PI, Sigma, USA) added to each sample. A maximum cell concentration of 500 000 cells/ml was accepted for each culture for staining. All processes must operate on ice and samples must also be maintained on ice until analysis by FCM. Histograms were analyzed using the internal software of FCM (BD FAC Station data processing system), determining peak position and the relative ploidy index of the samples (Galbraith *et al.,* 1983).

### 2.4. Analytical methods

Biomass was determined by filtering 20 mL of algal culture through a pre-weighed Whatman GF/C filter. The filter with algae was dried overnight at 60 °C in a hot air oven and weighted again to estimate the final dry weight. To obtain fresh weight, the Whatman filter was wetted with culture medium then weighed. After filtering 20 mL algal culture by vacuum pump, the wetted filter with fresh biomass was weighted again.

The amounts of the chlorophyll a, chlorophyll b, total chlorophyll and carotenoids were measured spectrophotometrically as described by Lichtenthaler, (1987).

Total soluble carbohydrates and starch were analyzed according to the methods employed by Roe (1955) and Thayumanavan and Sadasivam, (1984). Proteins were extracted according to Bradford (1976) spectrophotometrically.

To determine glycerol content, 4 mL of the growth culture of *D. salina* was centrifuged at 1500 rpm for 10 minutes at room temperature, and then the precipitated pellets were washed twice in a solution of 1.5 M NaCl and 5 mM phosphate buffer at pH 7.5. One mL of periodate reagent and 2.5 ml of acetylacetone reagent were added to 200 μl of precipitated pellets. The mixture was incubated at 45 ^o^C for 20 minutes. After that, optical density was determined at 410 nm. Results were compared with the prepared standard curve by using known amount of glycerol (Chitlaru and Pick, 1991).

The level of lipid peroxidation was determined by estimating malondialdehyde (MDA) and other aldehydes content using thiobarbituric acid (TBA) as the reactive material following the method employed by Heath and Packer, (1968).

### 2.5. Enzyme assay

Enzymes assayed in this work were extracted by grinding 0.15 g of fresh algae in a porcelain mortar contain 1.5 mL phosphate buffer containing 50 mM (pH 7.5) ethylene diamine tetraacetic acid (EDTA), 1 mM phenyl methanesulfonyl fluoride (PMSF), and polyvinylpyrrolidone (PVP) 1%. The extract was centrifuged for 15 minutes at 4 °C in 14000 g and the supernatant was assayed for enzymatic activity and quantification of protein by the Bradford method (1976). All operations were performed at 4°C. The activity of superoxide dismutase (SOD) (EC 1.15.1.1), catalase (CAT) (EC 1.11.1.6) and Guaiacol peroxidase (GPX) (EC 1.11.1.7) was determined according to the methods by Giannopolitis and Ries (1977), Azevedo *et al.* (1998) and Urbanek *et al.* (1991), respectively.

### 2.6. Statistical Analysis

The experiment was arranged in a completely randomized design with three replicates. SPSS software was employed for statistical analysis, and graphs were plotted by Excel software. Means were compared using Duncan’s multiple range tests at P<0.05.

## 3. Results

Result of figure 1, indicated that in the histogram a) the percentage of control cells were stopped in phase G_2_ to M is 30.63% and for the treated cells with 0.1% colchicines this percentage is relatively two fold, equals to 58.26% (Fig. b) which indicates that the cells are diploid. For the treated cells with 0.5% colchicine, percent of relative fluorescence intensity of nuclei (Fig. c) is 74.19%. In the histogram a) that is related to control, the peak in channel 160 is indicating the haploid cells and in two other histograms, channel 320 is indicating diploid cells and in both histograms also shows that there are still haploid cells but with increasing of colchicine concentration, the percentage of haploid cells have been lower. Therefore, the cultures treated by 0.1 and 0.5 % colchicine contained 27 and 43 % polyploid cells, respectively in comparison to control cultures.

**Figure 1.**
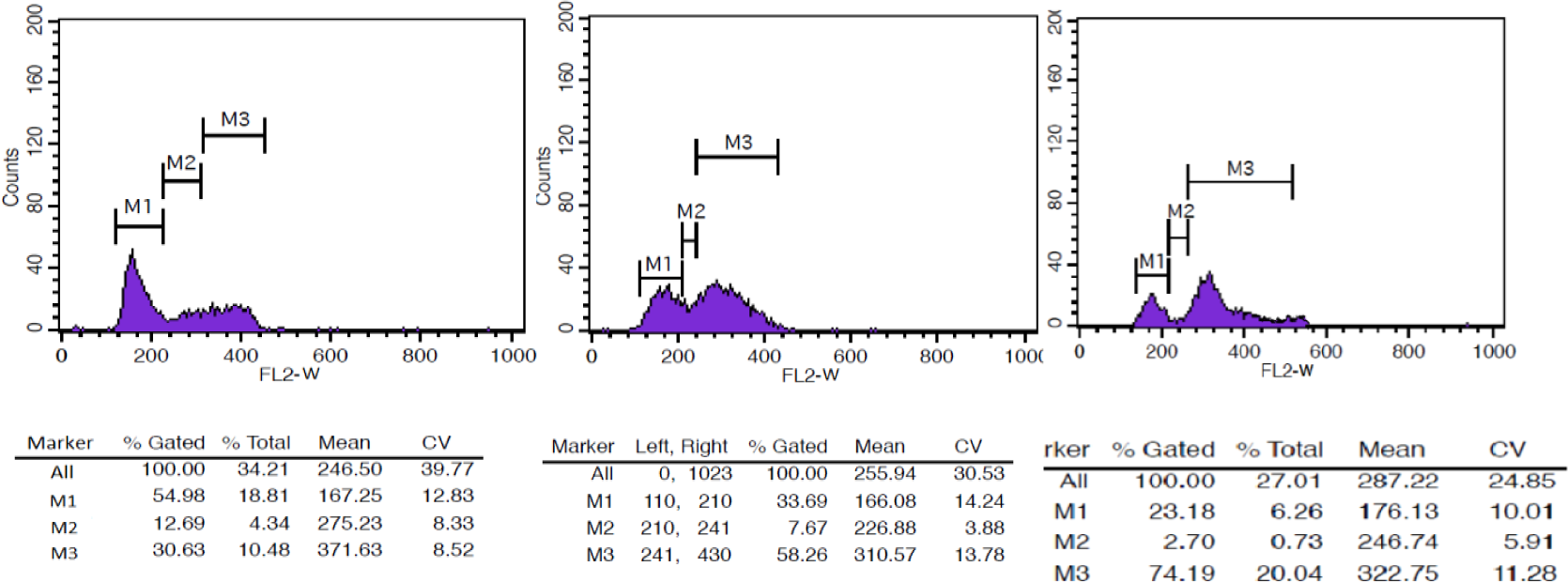
Histograms of relative DNA content of *D. salina* a) Histogram of relative fluorescence intensity of nuclei isolated from the control cells, b) the treated cells with 0.1% colchicine and c) the treated cells with 0.5% colchicine).

The results related to algal growth indicated that the highest fresh and dry weight of *D. salina* was observed in concentration of 3 M NaCl (Fig. 2). Algae exposed to 4 M NaCl contained the least amount of fresh and dry weight. In concentrations ranging from 2 to 4 M, it was found that 0.1 % colchicine had improvement effect on fresh and dry weight, respectively in the treated algae. Algae exposed to 0.5 % colchicine only showed an increase in dry weight in 3 M Nacl.

**Figure 2.**
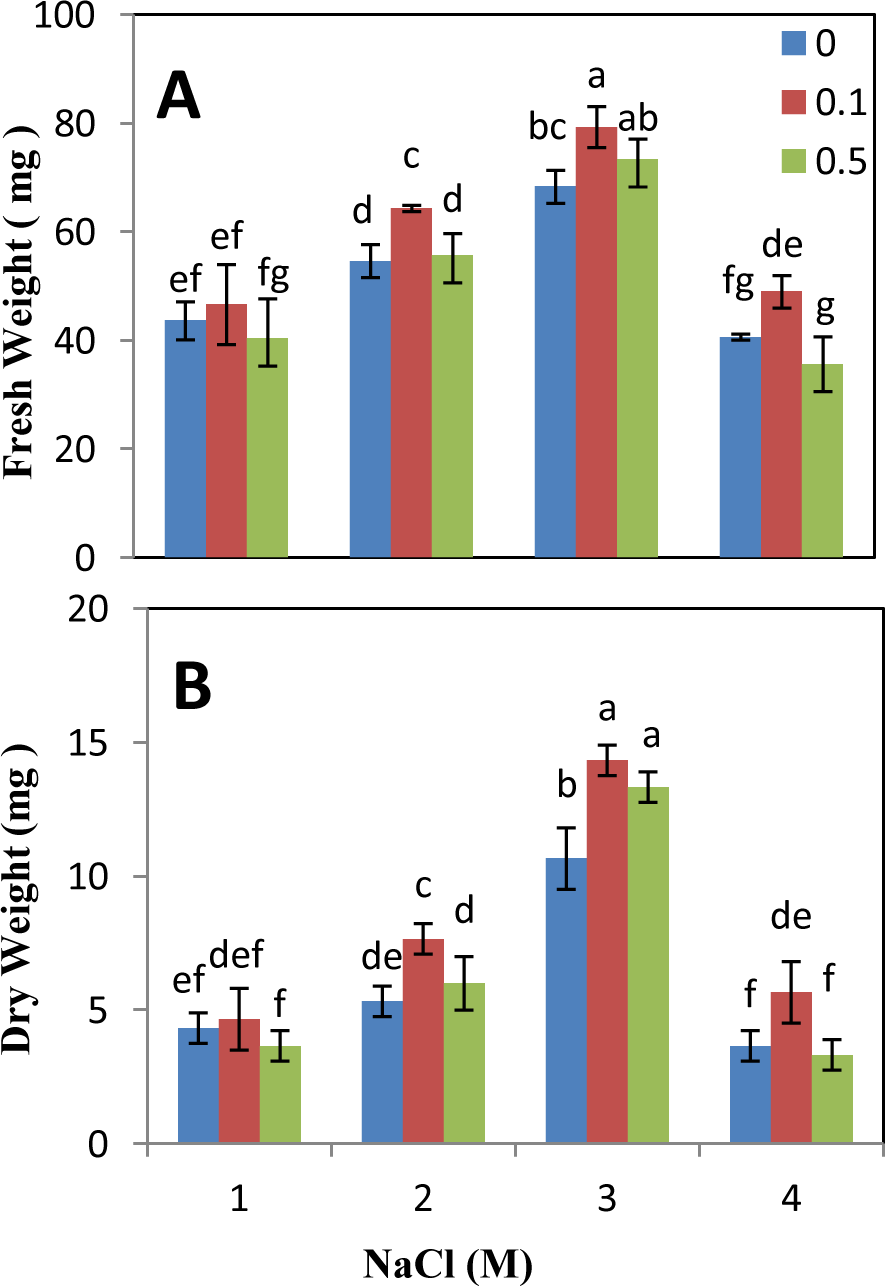
Effect of different concentrations of colchicine on fresh and dry weight of *D. salina* growing in media containing different concentrations of NaCl. The vertical bars represent the standard deviation; different letters above the bars indicate significant level at p < 0.05.

Fig 3 (A–D) demonstrates the impact of colchicine and salinity on chlorophyll and carotenoid content in *Dunaliella*. At higher and lower concentrations of 3M NaCl, a significant reduction in the amount of chlorophyll was observed. The lowest amount of chlorophyll was observed in 1 M salinity. In this respect, chlorophyll b was more sensitive than Chlorophyll a. Concentration 0.1% colchicine caused an enhancement in chlorophyll a and b content compared with control samples (without colchicine) at 2 and 3 M NaCl, and treatment of 0.5% colchicine had no significant change compared with controls to salinity 3 M, but at 4 M NaCl caused an increase in content of chlorophyll a and b.

**Figure 3.**
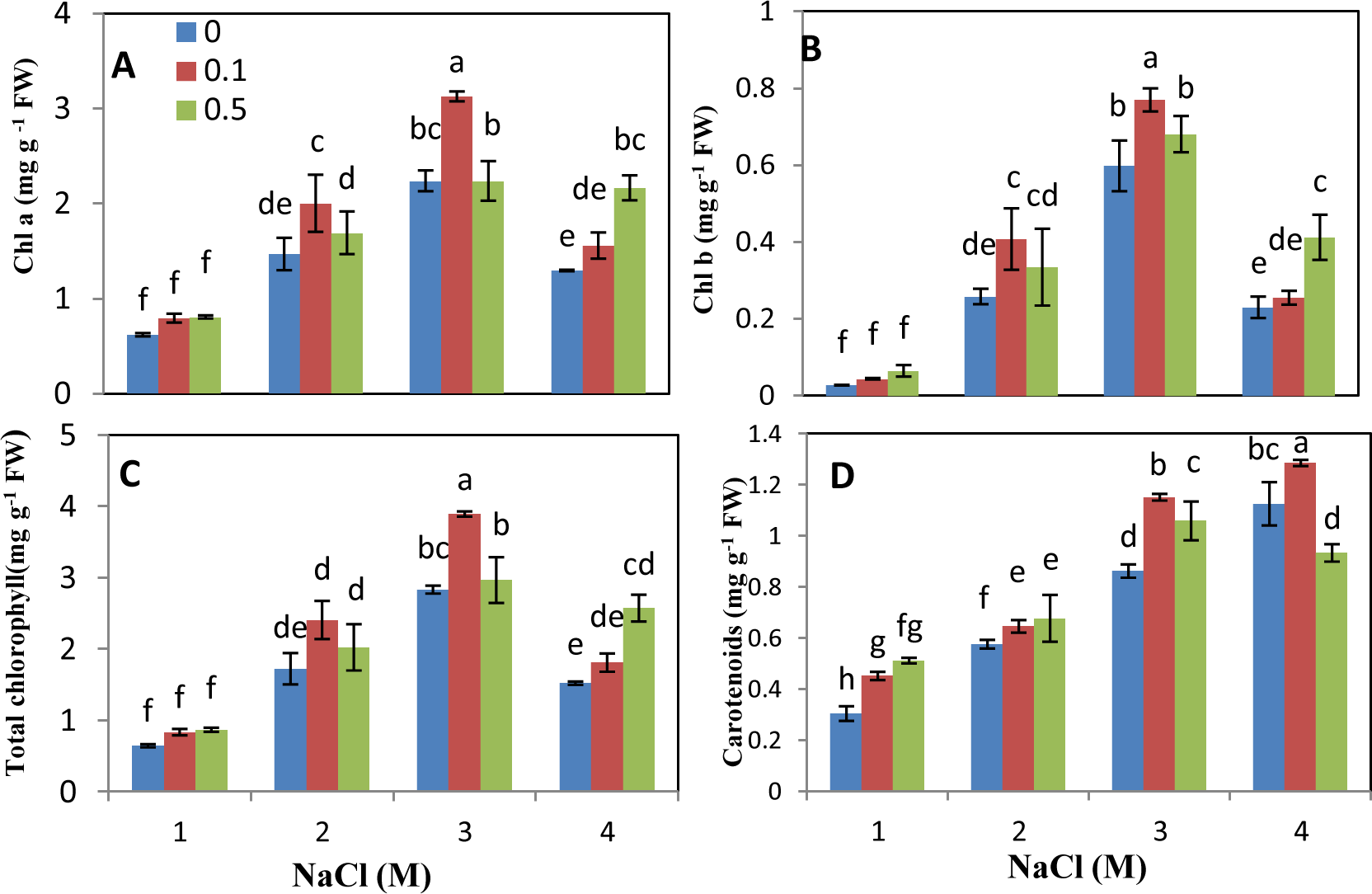
Effect of different concentrations of colchicine on (A) chlorophyll a and (B) chlorophyll b and total chlorophyll content (C) of *D. salina* growing in media containing different concentrations of NaCl. The vertical bars represent the standard deviation; different letters above the bars indicate significant level at p < 0.05.

Carotenoid content enhanced with increasing of salinity in culture. The highest cellular carotenoid level was recorded at 4 M NaCl. Algae exposed to 0.1% colchicine showed enhancement of carotenoids accumulation at all salinities (1- 4 M) (Fig. 3 D). Treatment of 0.5% colchicine increased the carotenoid content in the concentration range of 1-3 M NaCl.

The decrease in carbohydrate content (Fig. 4A) was observed under 1 and 4 M salinity. Treatment of 0.1% colchicine positively affected soluble sugar content under 3 and 4 M salinity. In addition, a significant increase in sugar content was noticed in the algae treated with 0.5 % colchicine and 3 M Nacl.

**Figure 4.**
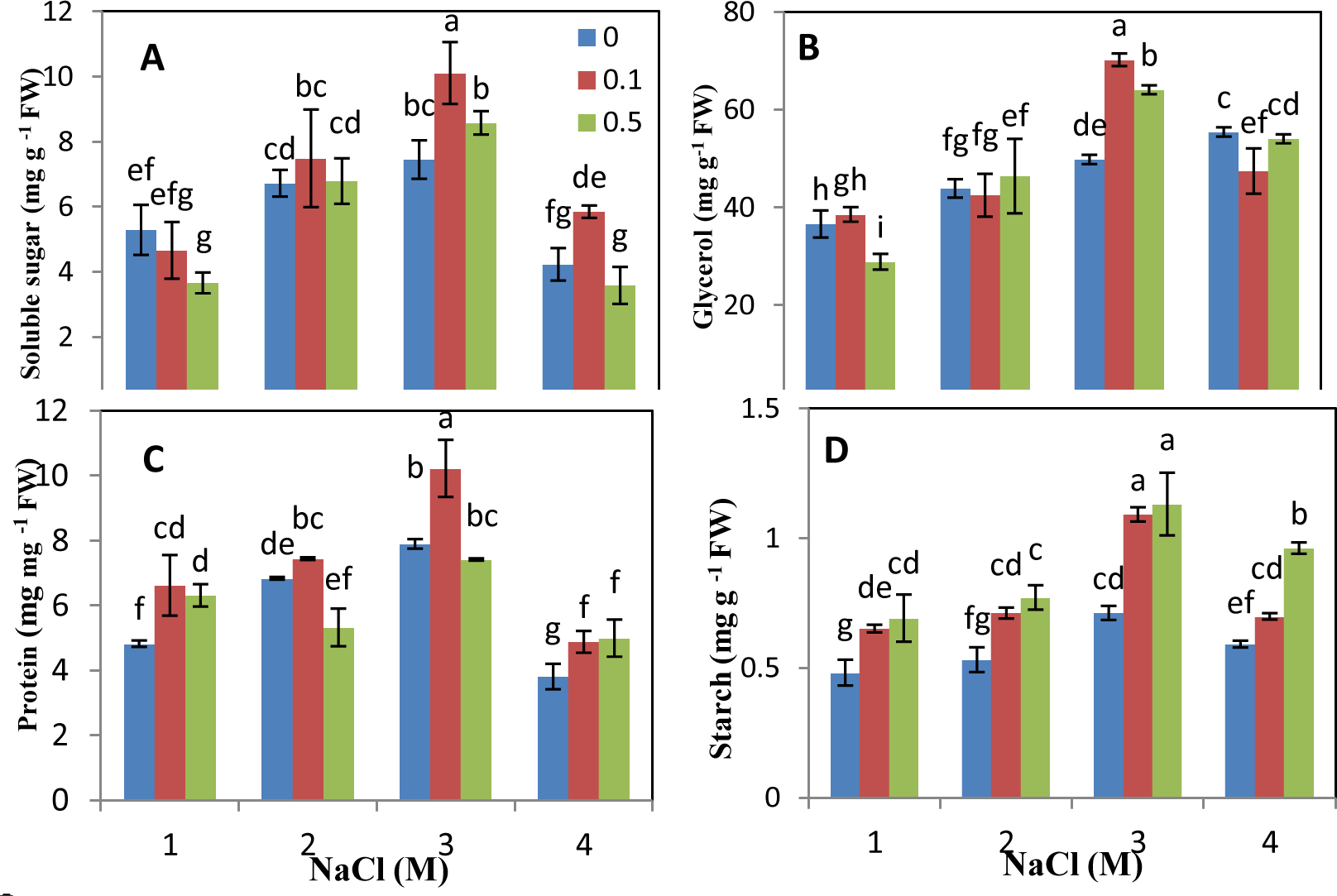
Effect of different concentrations of colchicine on (A) sluble sugar, (B) glycerol, (C) protein and (D) starch content (C) of *D. salina* growing in media containing different concentrations of NaCl. The vertical bars represent the standard deviation; different letters above the bars indicate significant level at p < 0.05.

After the exposure to different salinity levels, a considerable change in the glycerol concentrations was evident in untreated-colchicine samples (Fig. 4B). The maximum production of glycerol value was achieved in 4 M salinity. Induced, polyploidy created by 0.1 and 0.5 % colchicine considerably affected increasing of glycerol content at optimum salinity level (3 M NaCl).

Ranges of 1-3 M NaCl, stimulated protein accumulation (Fig. 4C). However, algae exposed to 4 M NaCl had the least amount of protein. The algae treated by 0.1 % colchicine showed an increase in protein content at all salinity levels. The highest increase was observed in the algae treated with 0.1 % colchicine and 3 M NaCl. Higher concentration of colchicine caused an increase in protein content under 1 and 4 M salinity.

The amount of starch was higher in the algae grown with 3 and 4 M NaCl (Fig. 4D). The algae treated by both concentrations of colchicine had more starch content at all salinity levels in comparison to untreated-colchicine algae. The highest increase was observed at 3 M NaCl with approximately 1.5 fold of untreated cells at this concentration (Figure 4D).

The present results showed a correlation between increasing of salinity level and that of MDA and other aldehydes content (Fig. 5). Algal cultures exposed to 0.1 and 0.5 % colchicine had less MDA and other aldehydes content under 2, 3 and 4 M NaCl. In this regard, concentration of 0.1 % colchicine was more effective. On the other hand, the results showed that lipid peroxidation was stimulated in the algae treated by 0.5 % colchicine under 1 M salinity.

**Figure 5.**
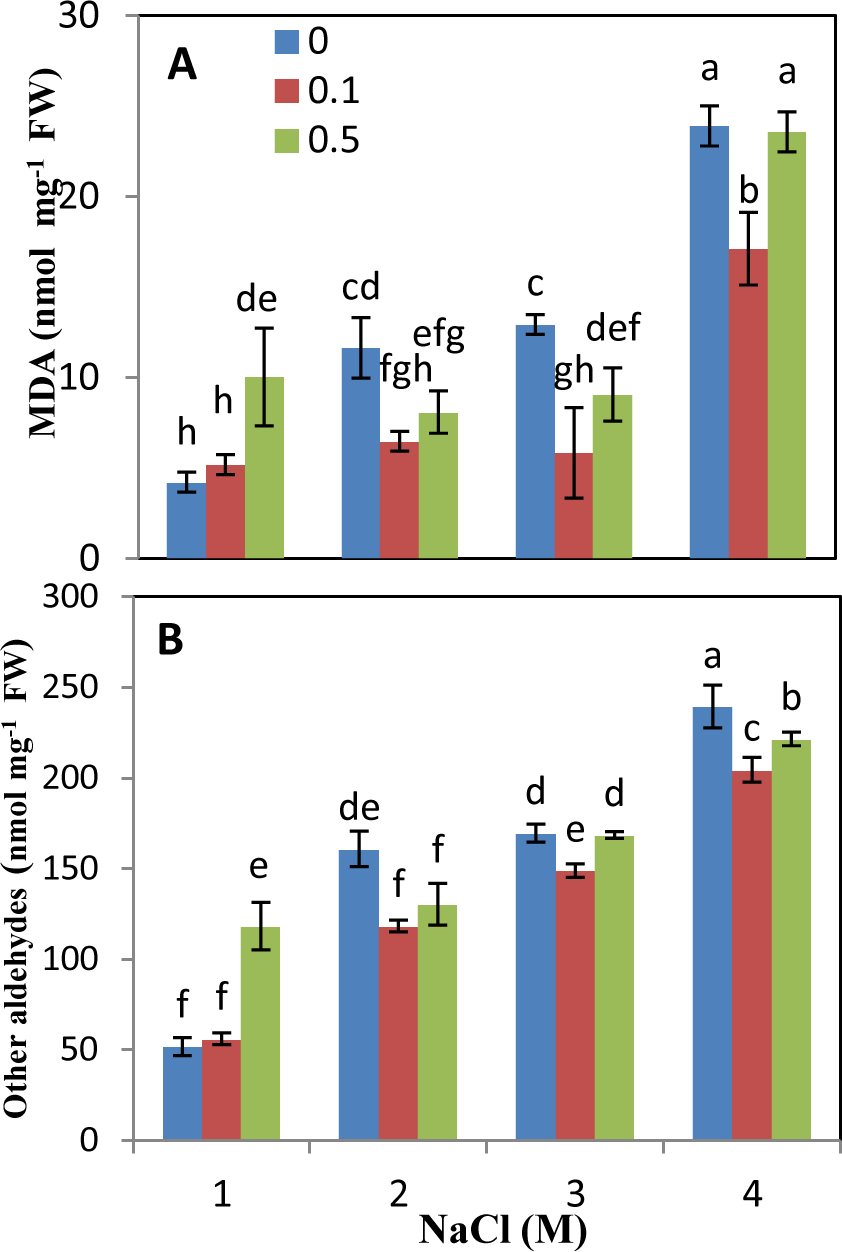
Effect of different concentrations of colchicine on MDA and other aldehydes content of *D. salina* growing in media containing different concentrations of NaCl. The vertical bars represent the standard deviation; different letters above the bars indicate significant level at p < 0.05.

Salinity significantly affected the activity of ROS scavenging enzymes (Figure 6A-C). Catalase activity in the algae grown under optimum salinity (1, 2 and 4 M) increased in comparison to cultures containing 3 M NaCl (Fig. 6A). Induced polyploidy by 0.1 and 0.5 % colchicine stimulated catalase activity at cultures containing 3 and 4 M NaCl. Polyploidy had no significant changes at two other salinity concentrations in catalase activity.

**Figure 6.**
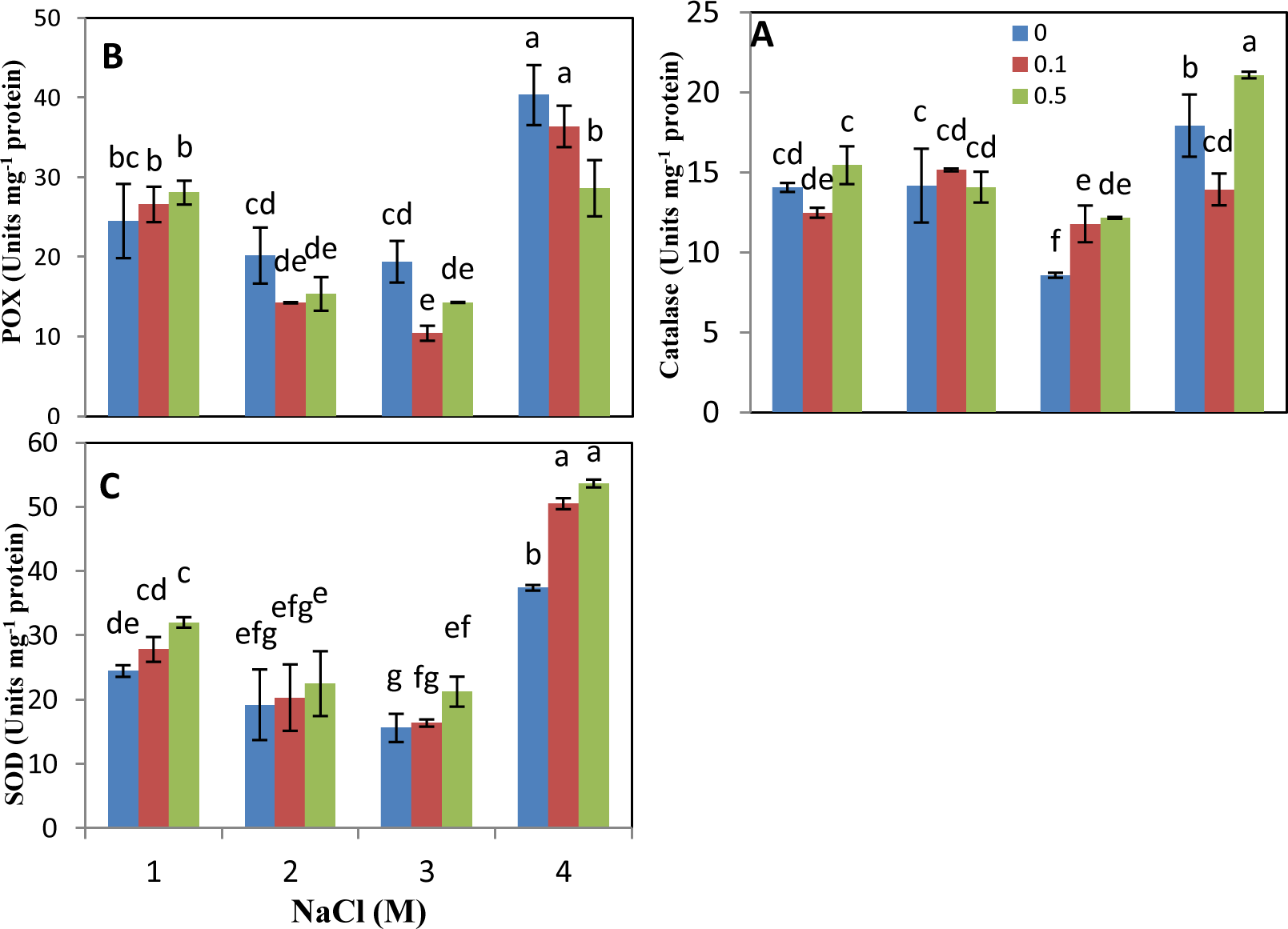
Effect of different concentrations of colchicine on (A) CAT, (B)POX and (C) SOD activity of *D. salina* growing in media containing different concentrations of NaCl. The vertical bars represent the standard deviation; different letters above the bars indicate significant level at p < 0.05.

The results revealed that the activity of peroxidase (PX) increased approximately 2 folds in the cultures treated by 4 M NaCl in comparison to other salinity treatments (1, 2 and 3 M NaCl) (Figure 6B). Specific peroxidase activity of treated algae by colchicine in salinities 3 and 4 M was lower than their controls. Like catalase, there was no significant change in the PX between control and colchicine treatments in the concentrations of 1 and 2 M NaCl.

The specific activity of SOD was almost doubled at 4 M NaCl in *D. salina.* Samples containing polyploidy cells showed remarkable increase compared to the control at the same concentration of salinity (Figure 6C).

## 3. DISCUSSION

Optimal concentration of NaCl to growth of *D. salina* was 3M, therefore the maximum fresh and dry weight content was gained at this concentration, but the fresh weight decreased with higher and lower concentrations of salinity. A negative trend in the growth rate of the microalga *Chlorella* sp. was observed when the NaCl concentration was further increased. This decrement in growth is due to the accumulation of reactive oxygen species (Kalita *et al.,* 2011). Cultures with 58 % polyploidy level (0.1% colchicine) had more fresh and dry weight at most salinity levels. However, higher percentage of polyploidy (74%) induced by 0.5% colchicine treatment increased dry weight only in the optimum condition (3 M NaCl). It suggests that presence of equal percentage from haploid and polyploid cells had improvement effect on the algae growth. In one experiment on diatom, the correlation between 4N polyploidy class (with 21% percentage of ploidy) and increased growth rate was showed. Vonshak and Richmond, (1981) have provided evidence that increase in the cellular DNA content in blue-green algae occurs in association with an increase in growth rate, indicating an increase in the number of genomes in the cell. D’Amato (2003) reported that the increased availability of nuclear templates was reflected in the growth of protoplast and its functional activity. He also noted the correlation between the rate of RNA synthesis and the cell ploidy level. Thus, increased availability of RNA templates allows higher level of protein translation and ultimately higher production of structural and functional proteins increasing the growth rate. Higher growth rate is desirable for commercialization as it allows more biomass to be produced within a shorter period of time. Bagheri and Mansouri (2015) reported that mixoploidy *Cannabis sativa* induced by colchicine treatment showed better growth compared to tetraploid plants. In spite of the importance of polyploidy, there are few studies on induction of polyploidy or its effect on algae, but in plants, it is well known that polyploidy leads to an increase in organ size, which may be caused by changes in activities of cell division and expansion as the result of the duplication of gene loci and increase in nuclear DNA content (Sugiyama, 2005). Furthermore, in other researches, plants with higher yield were observed by colchicine treatment compared with untreated plants (Mensah, 2007). Some varieties of potato responded significantly and positively in plant height, number of leaves formation and fresh weight of plant after colchicine treatment (Alam *et al.,* 2011).

The results showed a significant reduction in the amount of chlorophyll at higher and lower concentrations of 3 M NaCl, but in the colchicine treated cultures of *D. salina*, the amount of chlorophyll a and b significantly increased (especially in 0.1% colchicine). The total chlorophyll content of *Sesame indicum* L. treated with colchicine enhanced at higher concentrations. Contrary to these findings, with increasing ploidy level in *Juncus effusus*, chlorophyll content decreased significantly (Xu *et al.,* 2010).

There is a direct relationship between carotenoids content and increased levels of salinity. Since high salinity caused increment in carotenoids content and inversely low salinity caused decrease in carotenoids content. The photosynthetic apparatus does not use light energy sufficiently, and the excess energy leads to formation of free radicals rather than active oxygen molecules (singlet oxygen) under high salinity. These radicals are responsible for peroxidation reactions destroying diverse compounds of photosynthesis apparatus. Thus, the algae like *Dunaliella* and *Chlorella* accumulate large amounts of pigments for scavenging or omitting and reducing the radicals (ElBaz *et al.,* 2002). Induction of polyploidy in *D. salina* increased the carotenoids content, especially in 0.1% colchicine in all salinities. Comparative results of chlorophyll a, chlorophyll b, total chlorophyll, and carotenoids between tetraploid and diploid plants of cannabis demonstrated no significant changes (Bagheri and Mansouri, 2015).

The amount of protein and sugar in *Dunaliella* increased to 3 M NaCl, but in high salinity, their contents declined. However, Rao *et al.* (2007) reported a raise in carbohydrate content when the algae were cultured at different salinities. However, polyploidy raised protein and sugar content in some salt concentrations compared to their controls. Levine *et al.* (1990) in accordance with the present result reported that salt stress inhibited growth and development, decline in photosynthesis, respiration, and protein synthesis, and disturbed nucleic acid metabolism. Bagheri and Mansouri (2015) reported that tetraploid cannabis had higher values of total proteins compared with control plants (Bagheri and Mansouri, 2015). In the mentioned study, the highest amount of protein and sugar was found in mixoploid plants. Jaskani *et al.* (2005) reported that the total sugar content was comparable in diploid and tetraploid fruits of watermelon line. Like our results, Grange *et al.* (2003) observed higher total sugar content in triploid than in diploid fruit of watermelon, but in tetraploid cannabis, a reduction in carbohydrate content was observed due to decrease of CO_2_ fixation (Bagheri and Mansouri, 2015). It can be illustrated that in polyploid cells with higher DNA content, protein and sugar biosynthesis pathways were stimulated.

With increasing of salinity, the content of glycerol in this alga rose and reached its maximum in 4 M NaCl. The main task of glycerol in *Dunaliella* seems to be an osmotic regulator so that it is a compatible solute at high concentrations of salt protecting enzyme activity. According to Avron and Ben-Amotz (1992), glycerol is an important compound for osmotic regulation in *Dunaliella*. Polyploidy only in optimum growth condition, increased the amount of glycerol. These results showed that in spite of increasing in DNA content, the genes related to glycerol biosynthesis at high salinity level (4 M) are inactive.

In this study, the amount of starch increased at 3 and 4 M NaCl, but polyploidy induced starch accumulation at all concentrations of salinity. At the higher concentrations of salt in the medium of *D. parva* was observed the higher rate of glycerol synthesis whereas this relation does not exist between starch content and salinity of the medium (Gimmler and Moller, 1981). In polyploid races of *Atriplex confertifolia*, content of DNA, enzyme activity, photosynthesis per cell and cell volume all increased (Warner and Edwards, 1989). Higher photosynthetic rate in polyploidy samples can be explained by inducing starch accumulation. The leaves of mixoploid and tetraploid plants of *Cannabis sativa* L. compared to control plants contained higher amounts of starch (Bagheri and Mansouri, 2015).

Raja *et al.* (2007) maintained that part of the main physiological responses caused by stress conditions, including the flux of carbon between starch production in the chloroplast, synthesis of glycerol in the cytoplasm and accumulation of carotenoids. Goyal (2007) also observed that increased salt stress caused increment in the contribution of products of starch breakdown to glycerol synthesis. However, in the present study, we did not find a special relationship between these metabolites.

Results show that salinity significantly increased MDA and other aldehydes content in controls. However, in treated samples with 0.1 % colchicines, the amount of MDA declined considerably. MDA is an important indicator of the damage caused by salt stress (Verslues, 2006). Lower lipid peroxidation in polyploid algae can be attributed to protective effect of nonenzymatic antioxidant compounds (such as carotenoids) protecting the photosynthetic apparatus from damage by quenching triplet chlorophyll and singlet oxygen, in addition to the different strategies in the enzymatic system (Young and Frank, 1996). Salinity increased O_2_^-^ and H_2_O_2_ content in both diploid (2X) and tetraploid (4X) of *Robinia pseudoacacia*. However, 2X plants showed higher O2^-^ and H_2_O_2_ content compared with 4X plants at the end of 10 days of the experiment. Additionally, MDA content increased in 2X and 4X after salt treatment, and 2X showed much higher levels than those of 4X, indicating more damage to membranes in 2X plants (Wang *et al.,* 2013).

Gossett *et al.* (1996) suggested that protection from oxidative damage was induced under salt stress by more active ascorbate-glutathione cycles and a higher level of antioxidant enzymes such as CAT, SOD and POD. Accordingly, *D. salina* exposed to the high level of salinity (4 M NaCl) was significantly increased in SOD, CAT and GPX activity, indicating that these enzymes were utilized for enzymatically scavenging of AOS. Polyploidy had no remarkable impact on increase of antioxidant enzymes activity, except its impact on the SOD activity in the algae treated by 4 M NaCl.

Interestingly, the activities of the antioxidant enzymes (POD and APX) in 4× plants increased under salt stress. Similar to the present results, tetraploid *Robinia pseudoacacia* showed higher SOD activity than that of diploid plants, indicating that 4X had a more efficient enzymatic antioxidant system against salt stress and deal with ROS compared with 2X plant (Wang *et al.,* 2013).

In conclusion, we investigated the existence of different percentage of polyploidy cells in *D. salina* cultures and its response to salinity. As expected, all analyzed parameters showed variation in each of the three ploid populations. In general, mixoploid cultures showed better characterization than haploid cultures, especially in the optimum condition (3 M NaCl). Thus, populations with equal percent of haploid and polyploid cells (0.1 % colchicine treatment) had a higher function. We argue that polyploidy can be suitable to increase the algae productivity.

## Acknowledgements

This work was carried out in Shahid Bahonar University of Kerman, Iran. We also thank from Research Support Fund of Shahid Bahonar University.

